# SLECA: a single-cell atlas of systemic lupus erythematosus enabling rare cell discovery using graph transformer

**DOI:** 10.64898/2026.02.11.705246

**Authors:** Maoteng Duan, Yao Shi, Hao Tian, Qiuqin Wu, Xiaoying Wang, Bingqiang Liu

## Abstract

Systemic lupus erythematosus (SLE) is a highly heterogeneous autoimmune disease with complex immune and molecular dysregulation. While rare immune cell populations are increasingly recognized as critical drivers of disease pathogenesis and progression, the lack of sufficiently powered, comprehensive single-cell atlases has limited their systematic identification and characterization. To address this gap, we present SLECA, the first large-scale single-cell atlas of SLE, enabled by a novel graph-transformer framework for the interpretable discovery and analysis of disease-relevant rare cell populations. SLECA integrates 366 single-cell samples with standardized clinical and biological metadata, providing a comprehensive and analytically unified atlas of systemic lupus erythematosus. By enabling scalable integration and interpretable analysis, SLECA resolves 54 distinct cell types, including rare populations with critical disease relevance. Notably, we identify double-negative T cells (DNTs) as a disease-expanded population whose abundance correlates with clinical severity. Through in silico perturbation, we demonstrate that key transcription factors, specifically JUN and EGR1, can reprogram DNT cells toward conventional T-cell phenotypes, highlighting actionable regulatory vulnerabilities in SLE.

## Introduction

Systemic lupus erythematosus (SLE) is a prototypical autoimmune disease that affects approximately 3.4 million individuals worldwide^[1]^. It is characterized by a breakdown of immune tolerance, self-tolerance, persistent auto-reactivity, and chronic inflammation, leading to progressive multi-organ damage involving the skin, blood, joints, and brain. In severe cases, renal involvement manifests as lupus nephritis, a major cause of morbidity and mortality^[2]^. Despite decades of research, the biological basis of SLE remains incompletely understood, largely due to its profound clinical and molecular heterogeneity, which varies across patients and evolves over time, posing major challenges for diagnosis and treatment^[3]^.

Accumulating evidence indicates that this heterogeneity arises primarily from immune dysregulation at the cellular level^[4]^. In this process, rare immune subsets play crucial roles, such as double-negative T cells (DNTs) and plasmacytoid dendritic cells (pDCs) have been implicated as key drivers of immune imbalance and disease progression in SLE^[5, 6]^. Although their potential impact on the disease has been preliminarily identified, the specific characteristics, functions, and mechanisms of these rare populations in immune dysregulation remain poorly defined. A major barrier is the lack of sufficiently powered, systematic analyses capable of robustly identifying and characterizing rare cell states across heterogeneous patient cohorts.

Recent advances in single-cell RNA sequencing (scRNA-seq) have enabled unprecedented resolution of cellular heterogeneity in SLE, including the detection of rare and transitional cell populations and reconstruction of disease-associated regulatory programs^[7, 8]^. However, existing SLE single-cell RNA-seq datasets are fragmented across cohorts and platforms, limiting cross-study integration and comprehensive atlas construction. Moreover, widely used clustering approaches, such as Leiden^[9]^ and Louvain^[10]^, are sensitive to parameter choices and often fail to reliably detect rare populations in complex disease settings^[11]^. Although specialized rare-cell detection methods have been proposed, including FIRE^[12]^, GapClust^[13]^, TooManyCells^[14]^, GiniClust^[15]^,their robustness and scalability in highly heterogeneous autoimmune diseases remain limited. Although MarsGT^[7]^ performs strongly for rare-population detection, complementary single-cell ATAC-seq data remain scarce and difficult to match with existing datasets, limiting the broad application of mechanism-level integrative inference^[16]^.

To address these challenges, we introduce SLECA (Systemic Lupus Erythematosus Cell Atlas), the first comprehensive single-cell atlas of SLE. SLECA integrates all publicly available SLE single-cell RNA sequencing datasets, encompassing 366 samples and over 4 million cells from eight studies. Additionally, SLECA incorporates a novel single-cell graph-transformer framework, SarsGT (Single-cell RNA Analysis for Rare-population inference using a Single-cell Graph Transformer), as its core analytical tool, designed to enhance the identification and analysis of rare cell populations with improved accuracy and interpretability, overcoming the limitations of traditional methods. Through a unified analysis pipeline, we identified and annotated 54 distinct cell types, including three biologically significant rare populations: pDCs, CD8^+^MAIT cells, and DNT cells. Notably, DNT cells are significantly expanded in SLE patients, while almost absent in healthy individuals, highlighting their potential as biomarkers for disease activity. Further in vitro perturbation experiments revealed that knockout of key transcription factors EGR1 and JUN can reprogram DNT cells’ transcriptional states toward conventional T-cell phenotypes, shedding light on their critical role in immune regulation. Beyond cell annotation, SLECA offers multidimensional analytical features, including gene importance assessment, functional enrichment, cell-cell communication, and regulatory network modeling, providing a systematic and actionable framework for exploring the immune microenvironment of SLE. By establishing a unified and scalable single-cell atlas, SLECA fills a critical gap in SLE research and provides key data and methodological support for uncovering disease mechanisms, discovering novel biomarkers, and identifying therapeutic targets, with significant scientific value and clinical translation potential.

## Results

### Overview of SLECA

SLECA is designed to address the challenges of unstable rare cell identification and the difficulty in locating cell type-specific key genes within a shared state space in cross-cohort integration. We integrated SarsGT, a graph-transformer framework developed specifically for the interpretable discovery of rare cell populations, into SLECA. SarsGT represents scRNA-seq data as a cell-gene heterogeneous graph and trains a heterogeneous graph transformer using probabilistic subgraph sampling, specifically designed for rare cell identification. It jointly outputs posterior probability for cell clusters and gene attention scores at the cell level. This architecture enhances the visibility of low-abundance cellular signals while providing a biologically consistent and interpretable measure of gene importance aligned with cell-state variation (see **Figure 1A**; **Methods**).

**Figure 1.**
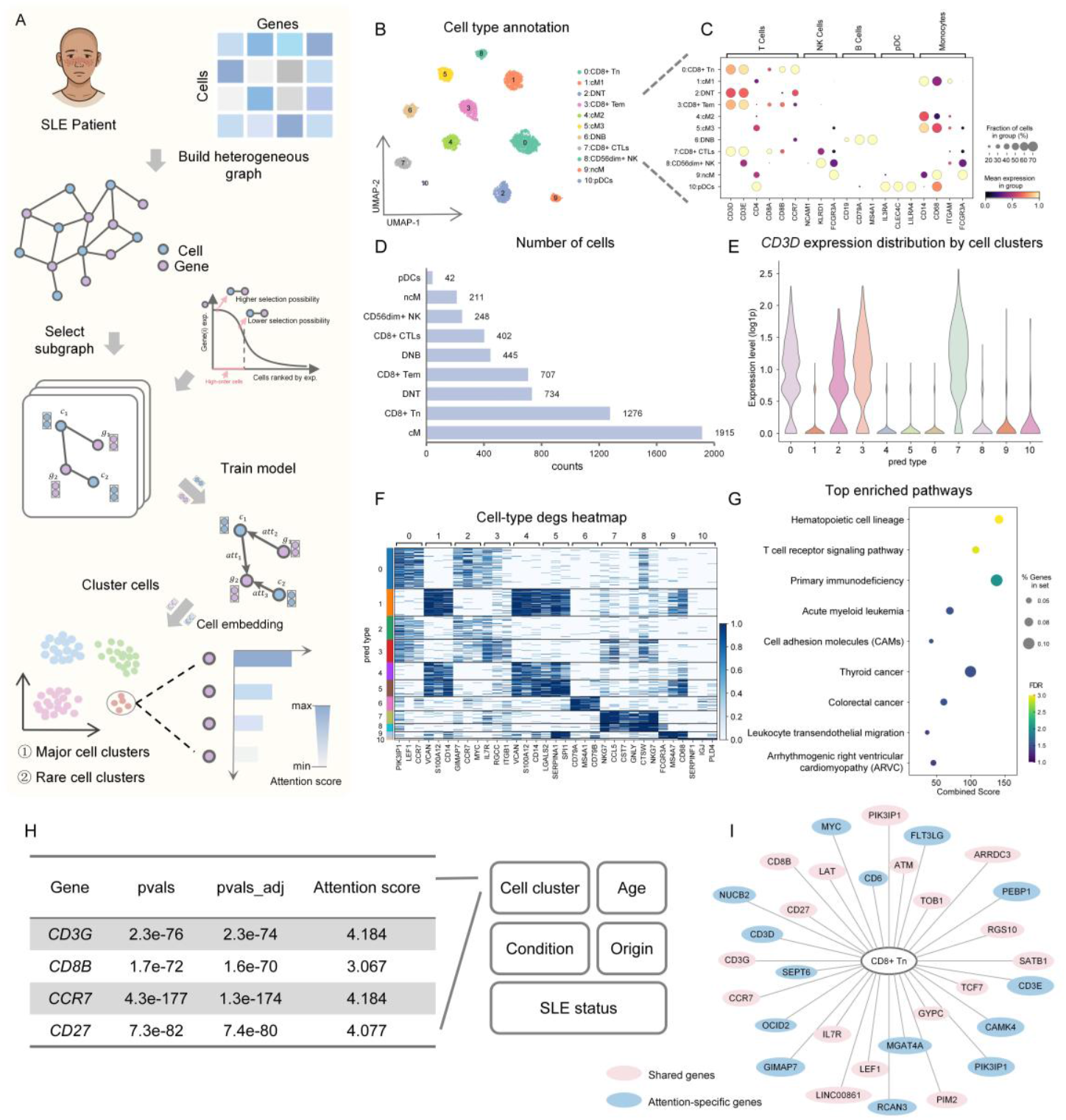
Overview of SLECA. **A)**, The framework of SarsGT. Features and visual representations related to scRNA-seq. **B**-**C)**, UMAP plot and dotplot of cell type annotations. **D)**, Barplot of cell counts across cell types. **E)**, Violin plots showing gene expression across cell types. **F)**, Heatmap of DEG among cell types. **G)**, Functional enrichment analysis of DEG. **H)**, Attention scores of different cell types. **I)**, Network visualization of genes associated with CD8^+^ Tn cells identified by the attention-based approach. Pink nodes represent genes shared with DEG analysis, whereas blue nodes are specific to the attention mechanism.

By leveraging SarsGT, SLECA performs a unified analysis across all samples, yielding multi-layered results. Within a shared embedding space, key immune subsets are resolved, including CD8^+^naive T cells (CD8^+^Tn), DNT cells, pDCs, and CD56^dim^ NK cells, thereby enabling cross-cohort comparisons (**Figure 1B-C**). The marker genes used for cell-type annotation are listed in **Supplementary Note N1 Marker gene**. Meanwhile, cluster compositions are quantified, and marker/representative genes are visualized, providing direct evidence for compositional differences across conditions and populations (**Figure 1D-E**). Examples of violin plots for additional genes are shown in **Supplementary Fig S1**. Moreover, SLECA includes differential expression genes and pathway enrichment results, for example, when comparing CD8^+^Tn cells with other subsets, immune-related pathways such as T-cell receptor signaling are significantly enriched (**Figure 1F-G**). Moreover, the attention mechanism of SarsGT assigns important scores to each gene in each SLECA cell and enables comparisons across cell clusters, age groups, disease states, tissue sources, and activity levels (**Figure 1H**). In comparison with traditional differential expression gene (DEG) analyses (**Figure 1I**), attention-based prioritization not only reproduces canonical marker genes but also captures additional regulatory genes closely associated with cellular phenotypes, such as *CD3D, CD3E*, and *MYC* highlighting its added value in discovering key genes.

### Cross-Cohort Data Integration in SLECA

SLECA compiles a total of 366 human scRNA-seq samples, derived from seven published SLE studies and one healthy control study (all obtained from the GEO database; sample details are provided in **Supplementary table S1-S9**). For each sample, we standardized key metadata including age, sex, and tissue of origin (**Figure 2A**).

**Figure 2.**
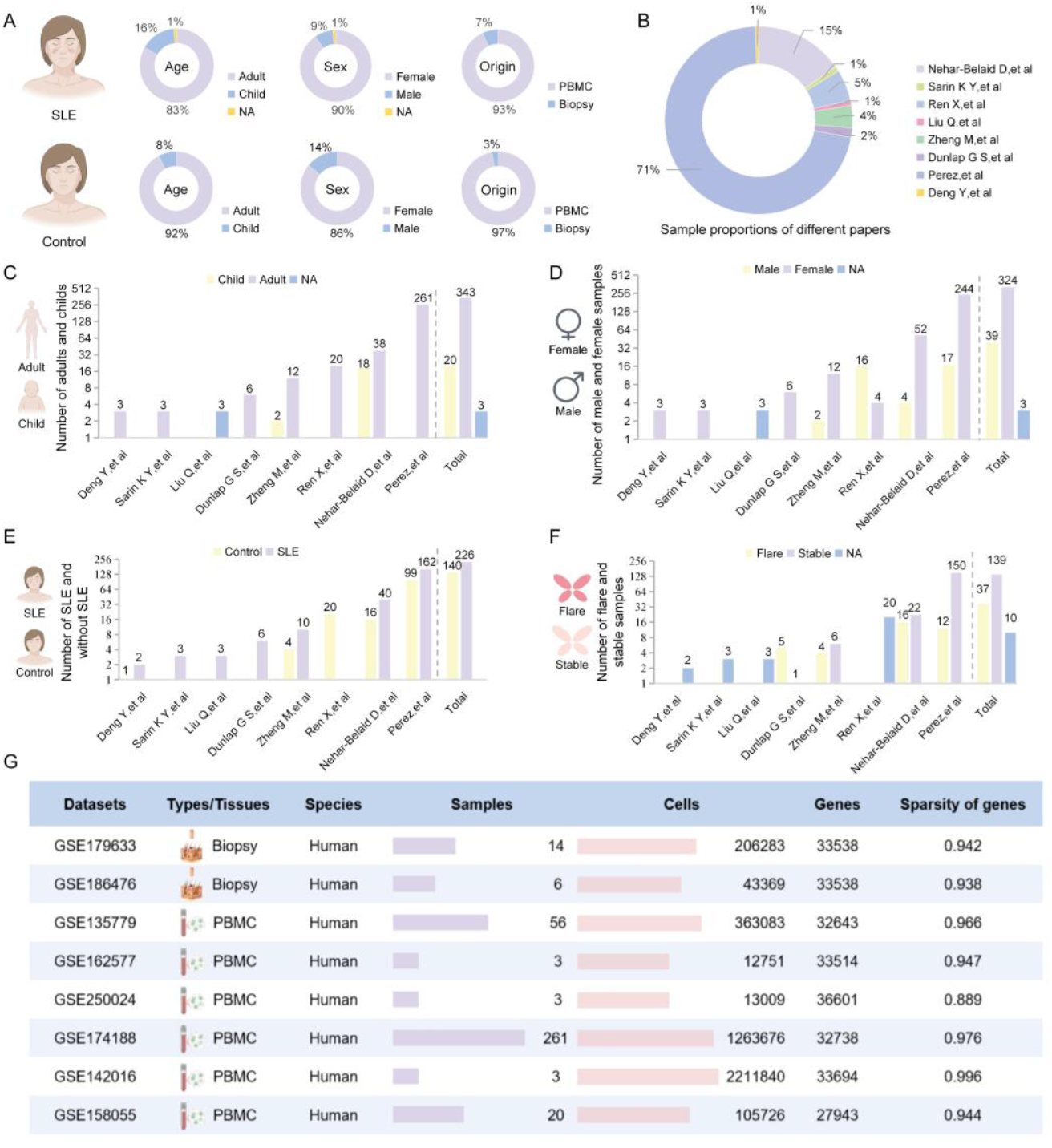
SLECA data characteristics and statistics. **A)** Proportions of samples by age, sex, and tissue source in within SLE and without SLE groups. **B)** Sample contributions across individual studies. **C-F)** Numbers and distributions of samples across studies stratified by (C) age, (D) sex, (E) disease status (SLE vs control), and (F) disease activity (flare vs stable). **G)** Dataset-level summary (GEO accession, tissue/source, species, numbers of samples/cells/genes, and gene sparsity).

Cross-study integration revealed that most samples originated from large-scale datasets, particularly Perez et al^[17]^, with the remaining studies serving as supplementary sources (**Figure 2B**). When stratified by age and sex (**Figure 2C-D**), the cohort was dominated by adult and female donors, consistent with the well-documented epidemiological pattern of SLE, which predominantly affects women of reproductive age. The atlas also includes both within SLE and without SLE samples (**Figure 2E**) and records disease activity states (flare/stable; **Figure 2F**), enabling case-control and activity-based comparative analyses. Furthermore, for each dataset, we systematically summarized accession numbers, tissue sources, and data characteristics, including sample, cell, and gene counts as well as data sparsity (**Figure 2G**), to facilitate methodological reproducibility and cross-cohort benchmarking. Detailed proportions and study contributions are provided in the figure legend and **Supplementary table S10-S11**.

### Cross-Condition Cell Type Composition and Distribution in SLECA

While cohort-level integration establishes the scope and representativeness of SLECA, its primary value lies in resolving disease-associated heterogeneity at the cellular level. Through SLECA, we annotated and constructed a comprehensive cell type landscape for both the within SLE and without SLE datasets, encompassing over 4 million cells and 54 distinct cell types. This analysis provides a foundation for a deeper understanding of the SLE immune microenvironment, highlighting significant differences in cell composition between the disease and control groups **(Figure 3A-B)**. These UMAP plots clearly reflect the spatial distribution of cell types in both the SLE and control groups, aiding in the identification of clustering patterns and distribution differences across the groups.

**Figure 3.**
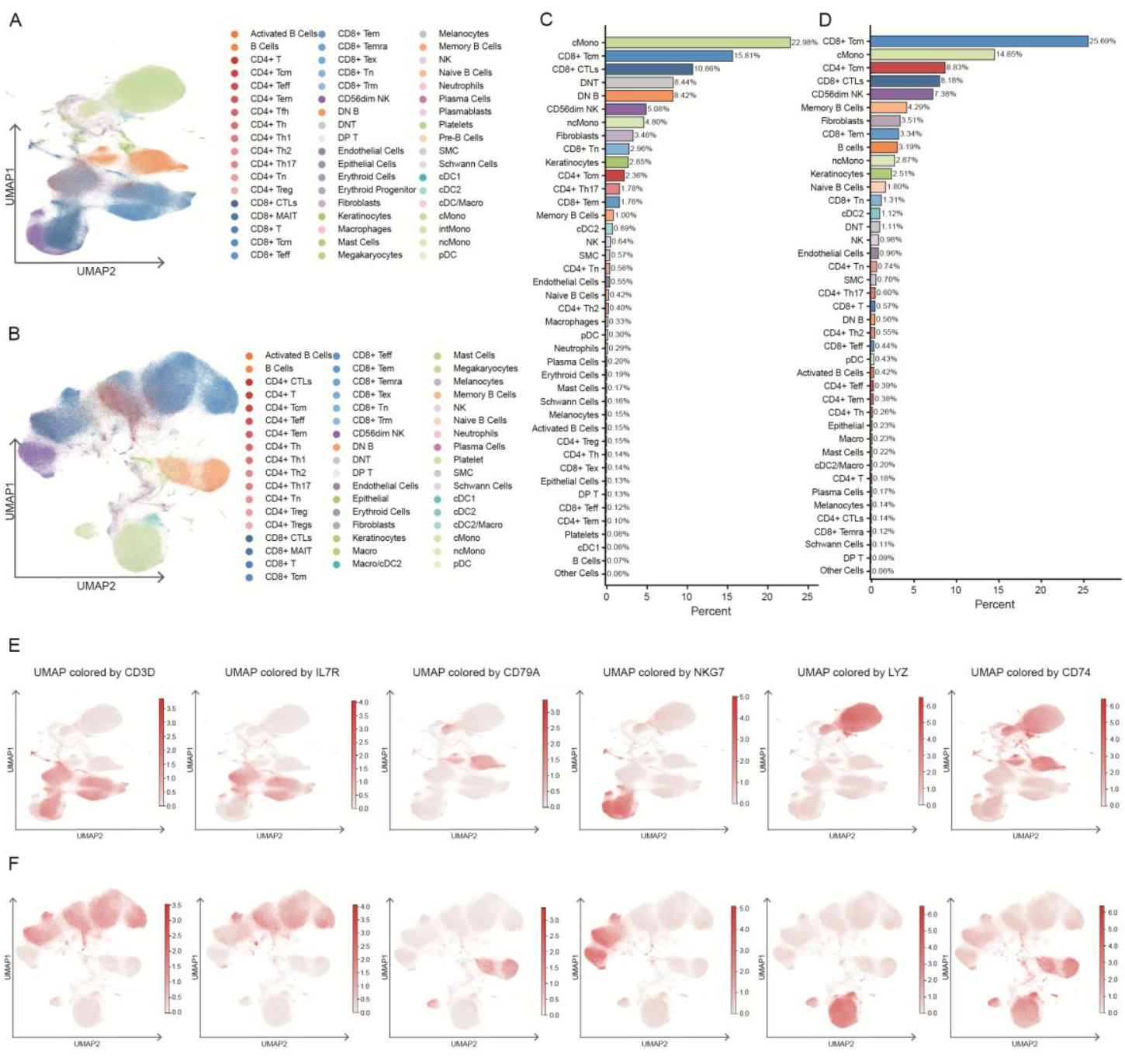
Comprehensive cell type landscape and marker gene expression in SLECA. **A-B)** UMAP visualizations of cell type distribution in SLE and control groups. **A**) SLE group, **B)** Control group. **C-D)** Bar charts showing the relative proportions of cell types in SLE and control groups. **C)** SLE group, **D)** Control group, highlighting variations in immune cell distribution (e.g., T cells, B cells, NK cells). **E-F)** UMAP visualizations of marker gene expression for major cell types (T cells, B cells, NK cells) in SLE and control groups. **E)** SLE group, **F)** Control group.

Cell type proportions are depicted in bar charts **(Figure 3C-D)**, illustrating the relative distribution of cell types in both the SLE and control groups. These plots emphasize the variation in the distribution of immune cells, such as T cells, B cells, and NK cells, across the groups. The detailed proportions of other cell types are provided in **Supplementary Note N2-N3**.

Additionally, we visualized the marker gene expression of major cell types, including T cells, B cells, and NK cells, in both the SLE and control groups. These visualizations reveal gene expression differences between the SLE and control groups **(Figure 3E-F)**. Other UMAP visualizations of marker gene expression are shown in **Supplementary Fig S2**. These multidimensional analyses of cell types and gene expression not only provide a detailed cell composition map of the SLE immune microenvironment but also lay the foundation for further in-depth studies of rare cell populations in SLE.

### Profiling DNT Cells in SLECA for Cross-Cohort Expansion and Functional Characterization

DNTs have drawn increasing attention in SLE due to their potential roles in pathogenic processes. These cells lack both CD4 and CD8 surface markers while expressing αβ or γδ T-cell receptors (TCRs), are extremely rare in healthy individuals but markedly expanded in SLE patients^[18]^. To investigate the role of DNTs in SLE, we focused on this cell population and found it to be associated with immune dysregulation and disease severity, consistent with previous findings^[6, 19]^.

We first examined their overall abundance and distribution in SLE and control groups. Specifically, samples within SLE and those without SLE were integrated separately to ensure within-group consistency. Based on cell clustering and annotation results from SarsGT, we integrated cell-type labels within each group and applied scVI^[20]^ for batch-effect correction. During correction, cell type was included as a covariate to minimize technical variation while preserving true biological differences. A group-specific unified UMAP was then generated to visualize the cellular landscape across samples. As shown in **Figure 4A**, DNT cells occupy a broader area with higher densities in SLE samples than in controls, providing clear evidence of their expansion in the disease context.

**Figure 4.**
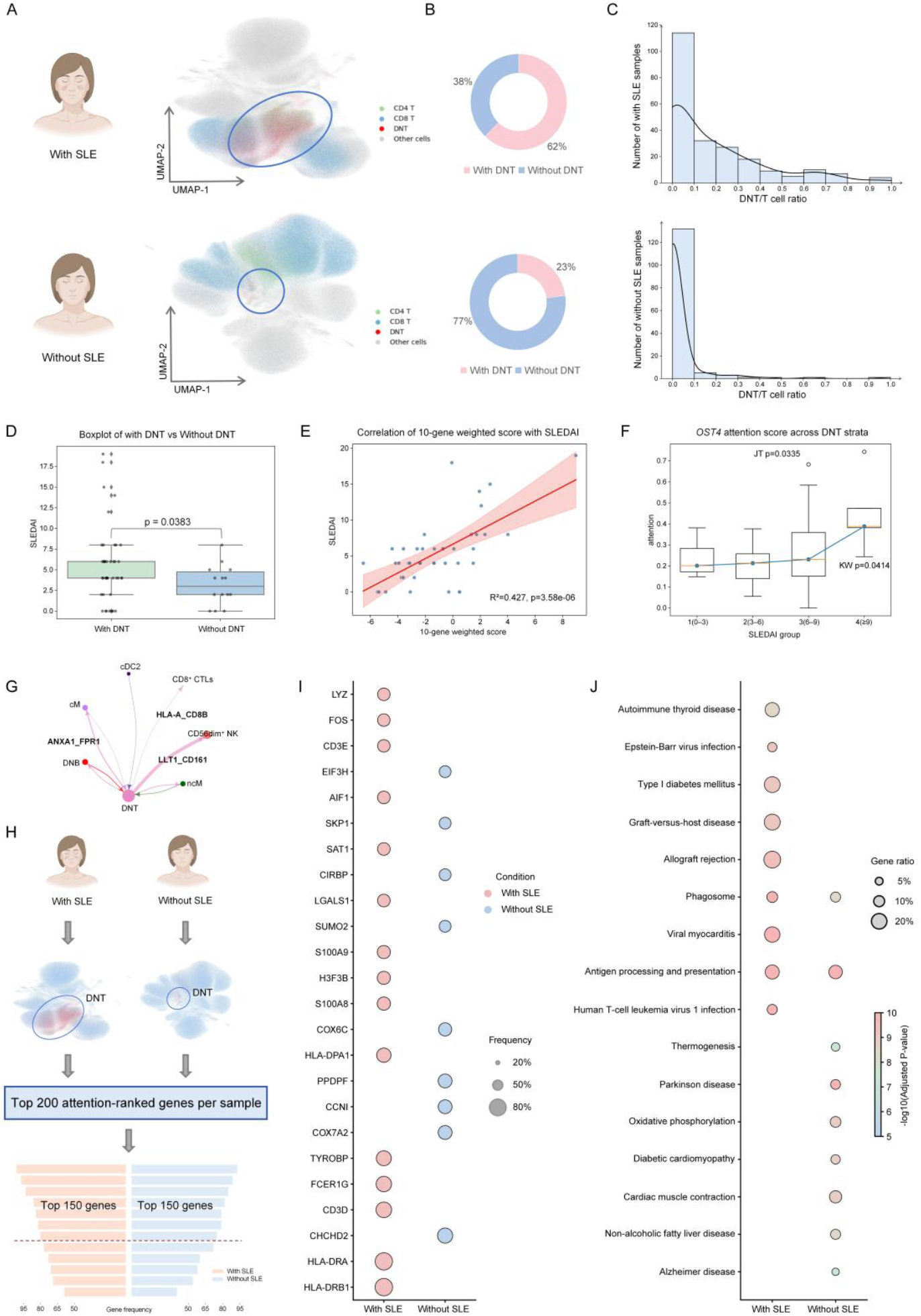
Functional characterization of DNT cells in SLE. **A)**, Visualization of DNT cell abundance, with DNT cells highlighted in red and other cell types in blue. **B)**, Proportion of samples within or without detectable DNT cells in SLE and control groups. **C)**, Distribution of the proportion of DNT cells among total T cells across samples, shown as a histogram with kernel density estimation (KDE). **D)**, Comparison of SLEDAI scores between samples within and without DNT cells, assessed using Mann-Whitney U test (*p* = 0.0383; *n*_1_ = 41, *n*_2_ = 14). **E)**, Attention-based quantification of gene importance in DNT cells. Multiple linear regression model constructed using the top 10 attention-ranked genes (*R*^2^ = 0.427, *P* < 3.58 × 10^−6^). **F)**, *OST4* attention scores across four SLEDAI strata (0-3, 3-6, 6-9, ≥ 9), showing a significant upward trend with disease activity (Kruskal-Wallis *p* = 0.0414 ; Jonckheer-Terpstra *p* = 0.0335). **G)**, Differential cell-cell communication networks between SLE and without SLE groups, highlighting DNT-centered signaling. **H)**, Workflow illustrates the selection of the top 150 key genes based on attention scores. The diagram outlines the stepwise procedure applied to DNT cells from both SLE and control groups, including gene ranking, frequency assessment, and final set determination. **I)**, Bubble plot showing the distribution of SLE-specific and control-specific genes in DNT cells, together with their occurrence frequencies. **J)**, Functional enrichment of DNT-specific genes in SLE.

We next compared the presence of DNT cells between the SLE and control groups at the sample level (**Figure 4B**). In the SLE group, 62% of samples contained detectable DNTs, compared to only 23% in controls, indicating a strong association between DNT occurrence and disease status. To quantitatively assess the degree of expansion, we calculated the fraction of DNTs among total T cells for each sample. The resulting distribution (**Figure 4C**) shows that control samples were mostly clustered near zero with few high-fraction cases, whereas SLE samples exhibited a right-shifted distribution with higher DNT/T ratios, consistent with DNT expansion and immune dysregulation in SLE.

To further evaluate the relationship between DNT expansion and disease activity, we analyzed SLEDAI clinical scores^[21]^ in SLE samples with available SLEDAI data (**Figure 4D**). Samples with detectable DNTs showed significantly higher SLEDAI scores than those without DNTs, supporting a positive association between DNT expansion and disease severity. This finding is consistent with previous studies reporting that elevated DNT frequencies are correlated with increased disease activity and tissue damage in patients with SLE^[22]^.

After establishing the expansion pattern and its clinical relevance, we leveraged the attention mechanism in SarsGT to prioritize genes and explore the potential mechanisms by which DNTs contribute to the pathogenesis of SLE. A multivariable linear model based on the top ten attention-ranked genes revealed a significant linear relationship with SLEDAI (*R*^2^ = 0.427, *P* < 3.58 × 10^−6^ ; **Figure 4E**), indicating that SarsGT captures key transcriptional regulators and functional genes closely linked to clinical activity. Regression plots for the individual gene-SLEDAI associations are shown in **Supplementary Fig S3**. We then stratified SLE samples by SLEDAI (0-3, 3-6, 6-9, ≥ 9) and examined attention scores across groups. *OST4* displayed a monotonic increase in attention with higher SLEDAI (**Figure 4F**), with the ≥ 9 group significantly exceeding lower groups, suggesting that *OST4* may participate in immune imbalance and disease progression and could represent a potential therapeutic target. Additional examples, including *ARPC3, MYL6*, and *SRP14* are shown in **Supplementary Fig S4**.

To further elucidate the molecular mechanisms underlying DNT-mediated immune dysregulation in SLE and to provide insights into potential restoration of immune balance, we compared intercellular communication patterns between the SLE and control groups. The differential network (**Figure 4G**) revealed that DNTs function as a communication hub in SLE, exhibiting markedly enhanced interactions with multiple immune subsets. Notably, LLT1-CD161 (CLEC2D-KLRB1) signaling between DNTs and CD56^dim^ NK cells was particularly prominent. Prior studies have shown that this signaling axis can co-stimulate T cells, enhancing their activation and cytokine secretion, while simultaneously suppressing NK-cell cytotoxicity^[23, 24]^. Consistent with these findings, our results suggest that DNTs may engage NK cells through the LLT1-CD161 pathway to form a “T-cell amplification and NK-suppression” feedback loop, thereby sustaining immune imbalance within the SLE microenvironment. In addition, HLA family ligand-receptor pairs between DNTs and other immune subsets were broadly strengthened, consistent with upregulated antigen presentation and T-cell activation, which may further promote aberrant B-cell activation and autoantibody production^[25]^. A more detailed bubble plot illustrating common DNT ligand-receptor interactions is provided in **Supplementary Fig S5** and **Supplementary table S12**. Building on the communication-level insights, we investigated the intrinsic molecular regulators of DNTs. Specifically, we integrated DNT cells from both SLE and control samples (**Figure 4H**). For each sample, the top 200 SarsGT attention-ranked genes were selected, and cross-sample occurrence frequency was calculated. To enhance the reliability of candidate genes, we further excluded genes with low occurrence frequency (below the first quartile). Representatives with SLE-specific and without SLE-specific genes among these candidates are shown in **Figure 4I**, and the complete list is provided in **Supplementary Fig S6** and **Supplementary table S13**.

To gain mechanistic insights into the biological functions of DNT-associated genes, we further pathway enrichment analysis using enrichR^[26]^(**Figure 4J**) revealed that DNT-associated genes were significantly enriched in Viral myocarditis, Antigen processing and presentation, and Epstein-Barr virus (EBV) infection pathways (FDR < 0.05). Enrichment in the first two pathways reflects activation of immune programs related to myocardial inflammation and enhanced antigen processing and presentation, both of which may contribute to the loss of immune tolerance and the maintenance of autoreactivity^[27, 28]^. Enrichment of the EBV infection pathway further supports the well-established role of EBV as an environmental trigger in SLE^[29]^. Although these findings provide correlative rather than causal evidence, when considered together with the intercellular communication results, they suggest that DNTs not only expand in SLE but also occupy a functionally central position interconnected across multiple immune pathways, potentially promoting disease activity through coordinated ligand-receptor signaling and key molecular programs.

### Identifying Disease-Associated Regulatory Modules and Potential Targets in the DNT Transcription-Factor Network within SLECA

Building on the gene and function level findings, we next probed transcriptional regulation underlying DNT biology. We applied ChEA3^[30]^ to the high-frequency gene set of DNTs derived from both SLE and control samples to predict upstream transcription factors (**Methods**). This analysis identified a restricted set of transcription factors significantly associated with the DNT-related genes (**Figure 5A**), suggesting regulation by a limited core module.

**Figure 5.**
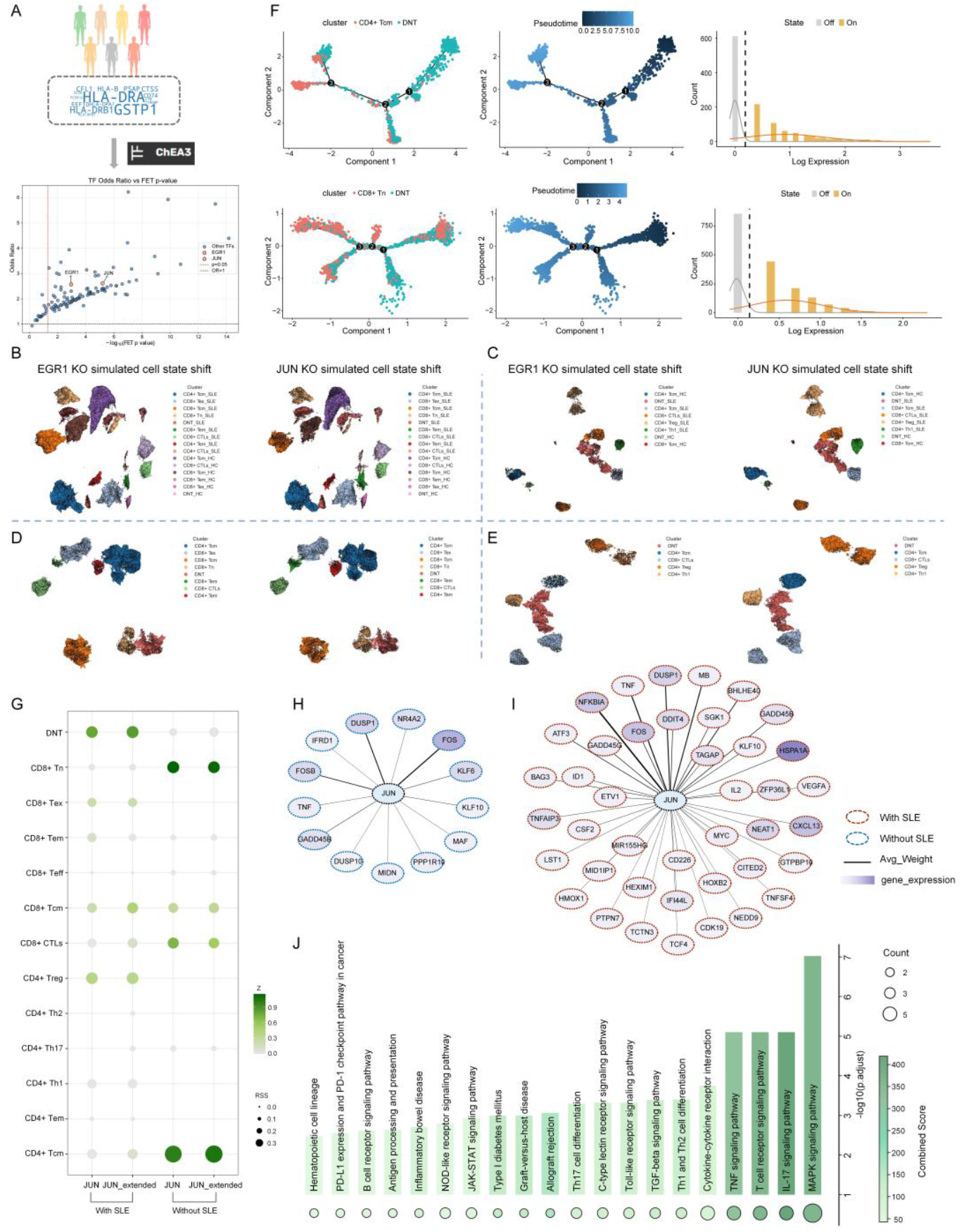
Functional characterization of JUN as a key transcription factor in DNT cells. **A)**, Transcription factor prediction based on high-frequency important genes identified from DNT cells in both within SLE and without SLE groups using CHEA3. The scatter plot displays enrichment results, with the x-axis representing FET significance (−log10 p value) and the y-axis representing enrichment strength (Odds Ratio). Key transcription factors EGR1 and JUN are highlighted, with the red dashed line indicating the significance threshold (*p* = 0.05) and the gray dashed line indicating OR = 1. **B)**, Extrapolated future states (arrows) after *JUN* knockout in PBMC-derived T cell subsets, integrating both within SLE and without SLE control samples. Colors indicate distinct T cell clusters. **C)**, Extrapolated future states (arrows) after *JUN* knockout in skin biopsy-derived T cell subsets, integrating both SLE and without SLE control samples. **D)**, Extrapolated future states (arrows) after JUN knockout in PBMC-derived T cell subsets from SLE patients only. **E)**, Extrapolated future states (arrows) after JUN knockout in skin biopsy–derived T cell subsets from SLE patients only. **F)**, Pseudotime trajectory of DNT cells inferred with Monocle2, along with visualization of *JUN* as a switch gene using Geneswitch. **G)**, Visualization of JUN regulatory activity across different T-cell subtypes in with SLE and without SLE samples. **H)**, Regulatory network as determined by SCENIC, based on JUN targets in without SLE samples. The network is relatively sparse with fewer downstream targets. **I)**, Regulatory network as determined by SCENIC, based on JUN targets in within SLE samples. The network shows increased density and higher average regulatory strength, with reinforced links to key targets (e.g., *NFKBIA, TNF, FOS, TNFAIP3*). **J)**, Visualization of functional enrichment results for JUN regulated genes in with SLE samples.

To evaluate the functional roles of these transcription factors, we performed knockout (KO) simulations using CellOracle^[31]^, focusing on JUN and EGR1 regulators (parameters and replicates are detailed in **Methods**). Prior studies have identified EGR1 as a potential regulator of DNT cell differentiation and function, in addition to its established roles in T-cell activation and inflammatory signaling^[32, 33]^. JUN, a central component of the AP-1 transcription factor complex, plays an essential role in T-cell activation, effector function, and lineage differentiation^[34, 35]^. Our simulations demonstrated that inhibition of JUN or EGR1 induced a reversal of DNT cell states toward conventional T-cell subsets under different data sources and experimental conditions. Specifically, in both the combined SLE-control dataset (**Figure 5B-C**) and the SLE-only dataset (**Figure 5D-E**), DNT cells exhibited a redistribution pattern approaching the regions corresponding to CD4^+^or CD8^+^T-cell subtypes. Consistent results obtained from additional datasets are shown in **Supplementary Fig S7-S9**. Pseudotime analysis with Monocle2^[36]^ further showed that 50% down-regulation of *JUN* expression redirected trajectories toward CD4^+^central memory T cells (CD4^+^Tcm) and CD8^+^Tn fates (**Figure 5F**). Complementary Geneswitch^[37]^ analysis suggested that *JUN* may act as a pivotal “switch” gene in determining DNT fate, where dysregulation may underlie DNT expansion and functional abnormalities in SLE. Analyses of additional datasets yielded concordant results (**Supplementary Fig S10-S14**).

To characterize JUN’s activity across T-cell subtypes, we applied SCENIC^[38]^ to SLE and control datasets (**Figure 5G**). Results showed that in SLE, the JUN (JUN_extended, representing regulons that include motifs linked to the TF by lower-confidence annotations such as those inferred by motif similarity) regulon exhibited the highest activity in DNT cells, whereas in controls, its activity was more pronounced in CD4^+^Tcm and CD8^+^Tn. This indicates that JUN activation undergoes cell-population-specific reprogramming under disease conditions, concentrating particularly in DNT cells.

We next contrasted JUN regulatory networks across conditions (**Figure 5H-I**). In controls, fewer targets were inferred, whereas in SLE samples, the number of JUN-regulated genes increased markedly. These downstream genes included *NFKBIA, TNF, FOS*, and *TNFAIP3*, all of which are tightly linked to reported SLE-associated SNPs, and many have been shown to be amplified or aberrantly activated in SLE patients^[39]^. For example, *TNF* exerts dual roles in SLE, functioning as both immunosuppressive and pro-inflammatory^[40]^; *NFKBIA* is a potential fundamental gene involved in the development of SLE, and its protein product, IκBα, influences the development and clinical manifestations of SLE^[41, 42]^. As another core component of the AP-1 transcription factor complex, *FOS* forms a heterodimer with JUN to constitute the AP-1 complex, which subsequently regulates JUN-associated transcriptional activity or cellular processes^[43]^. The *TNFAIP3* gene encodes the ubiquitin-modifying enzyme A20 and represents a well-established susceptibility locus for SLE, with its expression closely associated with disease activity^[44]^. These observations suggest that JUN and the genes constitute a disease-related transcriptional regulatory network in SLE.

Finally, functional enrichment of JUN-regulated genes (**Figure 5J**) revealed significant involvement in multiple pathways central to immune dysregulation and autoimmunity, including the TCR signaling, IL-17 signaling pathway, MAPK signaling pathway, TNF signaling pathway, and Antigen processing and presentation. Dysregulated TCR signaling is a hallmark of SLE, driving T-cell hyperactivation^[45]^; enrichment in IL-17 signaling and Th17 differentiation suggests JUN may exacerbate tissue damage via pro-inflammatory cytokine production^[46]^. Enrichment of MAPK and TNF signaling highlights JUN’s potential role in amplifying inflammation and cytokine network imbalance^[47, 48]^; while antigen processing and presentation underscores the breakdown of immune tolerance and persistent autoantigen stimulation^[28]^.

Together, these results demonstrate that JUN is aberrantly and specifically activated in DNT cells in SLE, with its regulatory network spanning multiple immune pathways central to disease pathology. JUN and its downstream targets (e.g., *TNF, NFKBIA, TNFAIP3*) not only align with known genetic susceptibility but also actively drive inflammatory and autoimmune processes, underscoring JUN as a key transcriptional regulator and a potential therapeutic target in SLE.

## Discussion

We developed SLECA, an integrated single-cell transcriptomic atlas for SLE that consolidates eight studies comprising 366 samples. The atlas incorporates an attention score derived from a graph transformer as a model-based, complementary, and non-causal indicator of gene importance alongside conventional DEG analysis (see **Methods**). Under a unified analytical framework, SLECA supports comparative analyses across age, sex, tissue source, and disease states.

As a representative finding, DNT cells showed significantly higher detection rates and relative proportions in SLE compared with controls. Moreover, JUN exhibited stronger regulatory activity within DNTs and was associated with multiple inflammation- and immunity-related pathways. Simulation analysis using CellOracle indicated that JUN inhibition could potentially drive DNT cells toward conventional T-cell states. While these findings provide correlational evidence that requires experimental validation, SLECA offers a consistent framework for cross-dataset reuse and validation, helping to robustly evaluate rare cell populations such as DNTs and generate mechanistic hypotheses.

The current version of SLECA has several limitations. It primarily focuses on transcriptomic data, with PBMC samples comprising the majority, while child, male, and tissue-biopsy samples remain underrepresented. Although scVI was applied for batch correction, residual batch effects may persist (see **Methods**). Future updates will incorporate spatial transcriptomics and multi-omics data to better characterize the tissue microenvironment of DNT cells and refine mechanistic inferences.

Overall, SLECA provides a reusable and interpretable single-cell resource for SLE, highlighting the potential importance of the DNT-JUN axis within the disease landscape and laying a foundation for future studies on diagnosis and therapeutic intervention.

## Methods

### Data Collection

We systematically searched and downloaded publicly available human scRNA-seq datasets related to SLE from the GEO database. A total of seven SLE studies comprising 346 samples and one control dataset including 20 healthy individuals were included (see **Supplementary Table S1-S9** for dataset details and metadata). For each sample, we uniformly annotated key metadata including age (child/adult), sex (male/female), tissue origin (PBMC, skin biopsy, etc.), disease status (SLE/control), study and platform identifiers, and other relevant variables to support stratified analyses and confounder control.

### Data Preprocessing

Preprocessing was performed using Scanpy v1.9.1^[49]^ and AnnData v0.8.0^[50]^: (1) Cell and gene filtering: Low-quality cells and lowly expressed genes were removed. The proportion of mitochondrial transcripts was calculated, and cells with abnormally high mitochondrial content were excluded. Thresholds for each dataset were adjusted based on a unified baseline and fine-tuned according to literature references and data distribution (sample counts are provided in **Supplementary Table S11**). (2) Doublet removal: Doublets were predicted and removed using Scrublet v0.2.3^[51]^, with the expected doublet rate estimated under default parameters. (3) Normalization and highly variable gene (HVG) selection: UMI counts were normalized to total counts per cell and log1p-transformed, followed by identification of HVGs. All quality control (QC) metrics were documented in the AnnData.obs and AnnData.var attributes to ensure traceability and reproducibility.

### Batch Correction

Cross-sample integration was performed using scVI v1.1.6^[20]^. The model incorporated batch as the primary batch variable and cell type as a categorical covariate to account for technical and biological variability. The resulting latent space was used for downstream visualization and statistical analyses. UMAP was applied solely for visualization. Key training parameters (latent dimension, number of epochs, learning rate, and random seed) and convergence diagnostics are provided in the **Supplementary Note N4 Model Parameters**.

### Rare Population-Friendly Clustering with SarsGT

To effectively identify rare cell populations, SLECA employed SarsGT for analysis. SarsGT is a single-cell clustering tool developed based on MarsGT^[7]^, our in-house tool for rare cell identification, which has demonstrated superior performance in rare cell detection. Unlike MarsGT, which requires paired multi-omics data, SarsGT relies solely on scRNA-seq data, effectively addressing the challenge of limited access to multi-omics samples. This tool retains the efficiency and interpretability of MarsGT in cross-dataset analysis, providing a robust solution for rare cell research under single-omics conditions. The core workflow of the SarsGT algorithm includes heterogeneous graph construction, subgraph sampling, feature learning and updating, model training, and cell cluster assignment.

### Heterogeneous Graph Construction

SarsGT begins with the raw scRNA-seq data, denoted as *X*^*R*^ ∈ ℝ^*M*×*N*^, where *M* is the number of genes and *N* is the number of cells. To capture the complex interactions between genes and cells, we construct a gene-cell heterogeneous graph denoted by *G* = (*V, E*), where *V* is the set of nodes and *E* represents the edges. Specifically, *V* = *V*^*C*^ ∪ *V*^*G*^, with 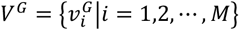, represents the set of gene nodes and 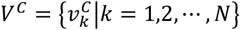 represents the set of cell nodes. The edges *E* in the graph are established between gene and cell nodes when 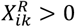, indicating a non-zero expression value for the gene in the respective cell. The initial feature representation for the nodes is derived from the raw data, denoted as *F* = *F*^*C*^ ∪ *F*^*G*^, where *F*^*C*^ = (*X*^*R*^)^*T*^ represents the feature matrix for cells, and *F*^*G*^ = *X*^*R*^ represents the feature matrix for genes.

### Subgraph Sampling

To enable efficient model training on large-scale datasets and enhance the detection of rare cell populations, SarsGT employs a subgraph sampling strategy before model training. This approach is motivated by the biological insight that genes with high expression in a small subset of cells play a pivotal role in both cell clustering and the identification of rare cell types. To construct the subgraphs for model training, SarsGT calculates a sampling probability for each gene-cell pair (*i, k*) as follows:

#### Step 1

For each gene *i* and cell *k*, an initial significance score *S*_*ik*_ is calculated. This score is defined as the ratio of the gene’s expression in the target cell *k* relative to its total expression across all cells:

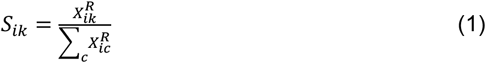

#### Step 2

The final sampling probability 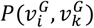 is derived through a conditional normalization process. For a given cell *k*_0_, genes whose expression exceeds a threshold *β* (defined as the first quartile of the gene expression value in *k*_0_) are selected, forming a high-expression gene subset 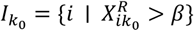. The sampling probability for gene *i*_0_ in cell *k*_0_ is then calculated as the ratio of its significance score 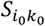 to the sum of significance scores for all genes in the subset 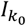:

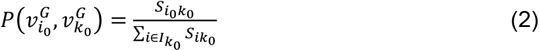

#### Step3

Based on the above probability, each training subgraph is constructed by randomly sampling 30 cells and sampling the associated gene nodes (i.e., neighbors) for each cell according to the computed probability distribution. To manage computational complexity, the number of neighboring nodes for each cell is limited to the smaller value between the total number of genes *N*_*G*_ and 20, i.e., min (*N*_*G*_, 20), ensuring that the subgraph size remains manageable.

### Feature Learning and Updating

The SarsGT model is trained on sampled subgraphs, using a stack of *L* layers of Heterogeneous Graph Transformer (HGT) to iteratively update node features. We denote the feature representation of node *v* at layer *l* as 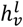. For the nodes *V*^*G*^ and *V*^*C*^, their initial features 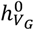 and 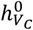 are directly derived from the initial feature matrices *F*^*G*^ and *F*^*C*^, respectively. To project different node types into the same latent space, we apply type-specific linear transformations *W* to the node features:

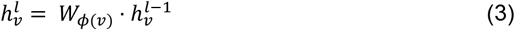

Here, *ϕ*(*v*) represents the type of node *v* (either gene or cell), and *W*_*ϕ*(*v*)_ is the corresponding learnable weight matrix.

Next, a multi-head attention mechanism is applied by dividing 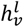 into *h* heads. For the *h*-*th* head at layer *l*, linear mapping functions are used: query (*Q*), key (*K*), and value (*V*). For each node *v*, the transformed representations are:

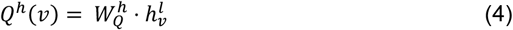

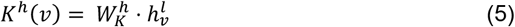

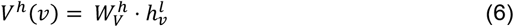

To compute the attention between node *v* and its neighbors *N*(*v*) in head h, we define an attention function estimating the importance of neighboring node *v*_*n*_:

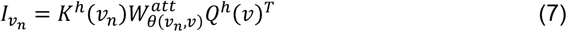

where 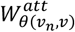 is a transformation matrix capturing edge type specific features, and *θ*(·) denotes the edge type. The overall multi-head attention is then computed as:

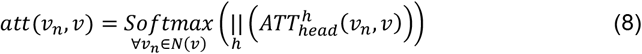

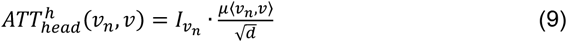

where ||(·) denotes the concatenation function, 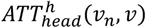 represents the attention weight of the *h*-*th* head, *μ* is a prior scaling function that accounts for the meta-relation characteristics between nodes and edges. Finally, the attention coefficients are normalized through the *Softmax* function to ensure that the attention weights between node pairs satisfy a valid probability distribution.

The information of node *v*_*n*_ in head *h* can be transmitted to node *v* through:

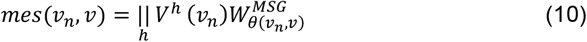

where 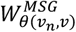 is a transformation matrix similar to 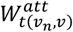.

To update the feature of node *v*, the final step at layer *l* aggregates the information obtained from its neighboring nodes *N*(*v*), based on 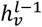 and the intermediate representation 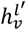:

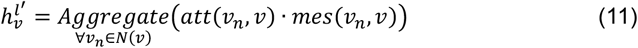

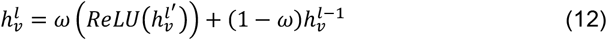

where *ω* is a trainable parameter and *ReLU* is the activation function. The aggregation function *Aggregate* can be implemented using average pooling, max pooling, or other pooling operations. The final embedding of the target node *v*_*n*_ is obtained by stacking the information from all *L* HGT layers. In the last HGT layer, the attention score between gene *i*_0_ and cell *k*_0_ is defined as:

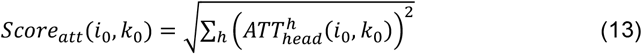

This score reflects the relative importance of gene *i*_0_ to cell *k*_0_. The formulation is designed to balance the contributions of positive and negative attention scores to the target node, ensuring that the model can fairly attend to information from different directions, thereby capturing the complex relationships between nodes more comprehensively. After multiple layers of updates, the final feature of node *v* is obtained.

### Model Optimization and Training

After the embeddings are calculated, the genes and cells obtain updated embeddings, denoted as 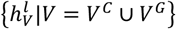. The update embeddings of cells 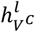 is denoted as *P* after normalizing by the sum of columns. Each row of matrix *P* represents a cell, each column corresponds to a manually defined reduced-dimensional set, and each element indicates the probability that a cell belongs to a specific cluster. In SarsGT, subgraph training is conducted in an unsupervised manner, with the loss function defined as follows: (1) Kullback-Leibler (KL) divergence loss, which preserves the structural consistency of the data. (2) Cosine similarity loss, which ensures that the features of cells within the same type are sufficiently similar. (3) Regularization loss, which promotes smoother optimization and faster convergence. The overall training objective is defined as:

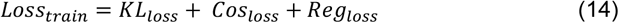

The KL divergence loss is formulated as:

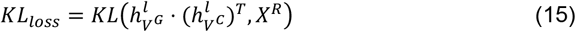

The Cosine similarity loss is defined as:

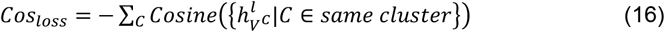

The regularization loss is defined as:

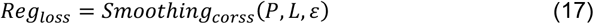

Here, *L* = {*l*_1_, *l*_2_, …, *l*_*N*_} represents the cluster labels obtained from Louvain clustering on the scRNA-seq data, and ε is a smoothing factor.

The smoothing cross-entropy loss is defined as:

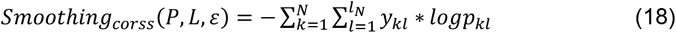

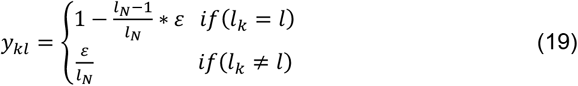

where *p*_*kl*_ represents the predicted probability of the given cell *k* belonging to the class *l* . These training objectives collectively help the model make confident predictions during optimization while maintaining stability, enabling SarsGT to learn robust cell-type representations across subgraphs.

### Cell Cluster Assignment and Cell Type Annotation

For each sample, SarsGT was applied to construct a cell-gene heterogeneous graph, followed by probabilistic subgraph sampling and message passing. The model output included a posterior probability matrix of cell clusters for each cell and cell-level gene attention scores. The attention score serves as a model-derived, interpretable indicator of gene importance, providing a complementary but non-causal measure alongside traditional DEG analysis. Cells were assigned to clusters based on the maximum posterior probability, and cluster identities were annotated using canonical marker genes (**Supplementary Note N1 Marker gene**). The results were visualized using dot plots, violin plots, and UMAP embeddings.

## Supplementary Information

### Authors’ contributions

Contributions: (I) Conception and design: Maoteng Duan, Yao Shi, Xiaoying Wang. (II) Data collection and programming: Yao Shi, Maoteng Duan. (III) Manuscript writing: Maoteng Duan, Yao Shi, Bingqiang Liu. (IV) Manuscript and figure revision: Qiuqin Wu, Hao Tian; (V) Final approval of manuscript: All authors.

### Funding

This work was supported by National Nature Science Foundation of China [NSFC, 62272270], and Shandong Provincial Natural Science Foundation for Distinguished Young Scholars [ZR2023JQ002].

### Data availability

All single-cell datasets used in this paper are publicly available. Data-1 was obtained from GEO under accession number GSE135779(https://www.ncbi.nlm.nih.gov/geo/query/acc.cgi?acc=GSE135779). Data-2 was obtained from GEO under accession number GSE142016(https://www.ncbi.nlm.nih.gov/geo/query/acc.cgi?acc=GSE142016). Data-3 was obtained from GEO under accession number GSE179 633(https://www.ncbi.nlm.nih.gov/geo/query/acc.cgi?acc=GSE179633). Data-4 was obtained from GEO under accession number GSE162577(https://www.ncbi.nlm.nih.gov/geo/query/acc.cgi?acc=GSE162577). Data-5 was obtained from GEO under accession number GSE186476(https://www.ncbi.nlm.nih.gov/geo/query/acc.cgi?acc=GSE186476). Data-6 was obtained from GEO under accession number GSE1 74188(https://www.ncbi.nlm.nih.gov/geo/query/acc.cgi?acc=GSE174188). Data-7 w as obtained from GEO under accession number GSE250024(https://www.ncbi.nlm.nih.gov/geo/query/acc.cgi?acc=GSE250024). Data-8 was obtained from GEO under accession number GSE158055(https://www.ncbi.nlm.nih.gov/geo/query/acc.cgi?acc=GSE158055). The processed data, including metadata, annotation information, embeddings, and attention scores, as well as the UMAP and dotplot visualizations, are available at the following link: https://zenodo.org/uploads/17698085.

### Code availability

The source code in this paper can be found at https://github.com/Snnrriet/SLECA.

## Acknowledgments

The authors thank Xue Liu and Xiaoyu Zhao for their discussions and suggestions.

## References

1. Siegel, C.H. and L.R. Sammaritano, Systemic lupus erythematosus: a review. Jama, 2024. 331(17): p. 1480–1491.

2. Zheng, M., et al., A single-cell map of peripheral alterations after FMT treatment in patients with systemic lupus erythematosus. Journal of Autoimmunity, 2023. 135: p. 102989.

3. Fanouriakis, A., et al., Update on the diagnosis and management of systemic lupus erythematosus. Annals of the rheumatic diseases, 2021. 80(1): p. 14–25.

4. Katsuyama, T., G.C. Tsokos, and V.R. Moulton, Aberrant T cell signaling and subsets in systemic lupus erythematosus. Frontiers in immunology, 2018. 9: p. 1088.

5. Liu, J., X. Zhang, and X. Cao, Dendritic cells in systemic lupus erythematosus: from pathogenesis to therapeutic applications. Journal of autoimmunity, 2022. 132: p. 102856.

6. Li, H., et al., Abnormalities of T cells in systemic lupus erythematosus: new insights in pathogenesis and therapeutic strategies. Journal of autoimmunity, 2022. 132: p. 102870.

7. Wang, X., et al., MarsGT: Multi-omics analysis for rare population inference using single-cell graph transformer. Nature Communications, 2024. 15(1): p. 338.

8. Chen, G., B. Ning, and T. Shi, Single-cell RNA-seq technologies and related computational data analysis. Frontiers in genetics, 2019. 10: p. 317.

9. Traag, V.A., L. Waltman, and N.J. Van Eck, From Louvain to Leiden: guaranteeing well-connected communities. Scientific reports, 2019. 9(1): p. 1–12.

10. Guillaume, L., Fast unfolding of communities in large networks. Journal Statistical Mechanics: Theory and Experiment, 2008. 10: p. P1008.

11. Grabski, I.N., K. Street, and R.A. Irizarry, Significance analysis for clustering with single-cell RNA-sequencing data. Nature Methods, 2023. 20(8): p. 1196–1202.

12. Jindal, A., et al., Discovery of rare cells from voluminous single cell expression data. Nature communications, 2018. 9(1): p. 4719.

13. Fa, B., et al., GapClust is a light-weight approach distinguishing rare cells from voluminous single cell expression profiles. Nature Communications, 2021. 12(1): p. 4197.

14. Schwartz, G.W., et al., TooManyCells identifies and visualizes relationships of single-cell clades. Nature methods, 2020. 17(4): p. 405–413.

15. Jiang, L., et al., GiniClust: detecting rare cell types from single-cell gene expression data with Gini index. Genome biology, 2016. 17(1): p. 144.

16. Yuan, Q. and Z. Duren, Continuous lifelong learning for modeling of gene regulation from single cell multiome data by leveraging atlas-scale external data. bioRxiv, 2023.

17. Perez, R.K., et al., Single-cell RNA-seq reveals cell type–specific molecular and genetic associations to lupus. Science, 2022. 376(6589): p. eabf1970.

18. Poddighe, D., et al., Double-negative T (DNT) cells in patients with systemic lupus erythematosus. Biomedicines, 2024. 12(1): p. 166.

19. Wu, Z., et al., CD3+ CD4–CD8–(Double-Negative) T cells in inflammation, immune disorders and cancer, Front. Immunol., 13, 816005. 2022.

20. Lopez, R., et al., Deep generative modeling for single-cell transcriptomics. Nature methods, 2018. 15(12): p. 1053–1058.

21. Bombardier, C., et al., Derivation of the SLEDAI. A disease activity index for lupus patients. Arthritis & Rheumatism: Official Journal of the American College of Rheumatology, 1992. 35(6): p. 630–640.

22. Dai, X., Y. Fan, and X. Zhao, Systemic lupus erythematosus: updated insights on the pathogenesis, diagnosis, prevention and therapeutics. Signal transduction and targeted therapy, 2025. 10(1): p. 102.

23. Tong, B., Immunobiology roles of the human CD161 receptor in T cells. Frontiers in Immunology, 2025. 16: p. 1648305.

24. Zhao, P., et al., The proportion of CD161 on CD56+ NK cells in peripheral circulation associates with clinical features and disease activity of primary Sjögren’s syndrome. Immunity, Inflammation and Disease, 2024. 12(4): p. e1244.

25. Jucaud, V., et al., Serum antibodies to human leucocyte antigen (HLA)-E, HLA-F and HLA-G in patients with systemic lupus erythematosus (SLE) during disease flares: Clinical relevance of HLA-F autoantibodies. Clinical & Experimental Immunology, 2016. 183(3): p. 326–340.

26. Kuleshov, M.V., et al., Enrichr: a comprehensive gene set enrichment analysis web server 2016 update. Nucleic acids research, 2016. 44(W1): p. W90–W97.

27. Saraca, L.M., et al., Cytomegalovirus myocarditis in a patient with systemic lupus erythematosus (SLE) successfully treated with ganciclovir. IDCases, 2018. 12: p. 4–6.

28. Xie, T., et al., Proteomics analysis of lysine crotonylation and 2-hydroxyisobutyrylation reveals significant features of systemic lupus erythematosus. Clinical Rheumatology, 2022. 41(12): p. 3851–3858.

29. Poole, B.D., et al., Aberrant Epstein–Barr viral infection in systemic lupus erythematosus. Autoimmunity reviews, 2009. 8(4): p. 337–342.

30. Keenan, A.B., et al., ChEA3: transcription factor enrichment analysis by orthogonal omics integration. Nucleic acids research, 2019. 47(W1): p. W212–W224.

31. Kamimoto, K., et al., Dissecting cell identity via network inference and in silico gene perturbation. Nature, 2023. 614(7949): p. 742–751.

32. Zhu, H.-R., et al., Double-negative T cells with a distinct transcriptomic profile are abundant in the peripheral blood of patients with breast cancer. Breast Cancer Research and Treatment, 2025. 209(1): p. 103–115.

33. Lohoff, M., et al., Early growth response protein-1 (Egr-1) is preferentially expressed in T helper type 2 (Th2) cells and is involved in acute transcription of the Th2 cytokine interleukin-4. Journal of Biological Chemistry, 2010. 285(3): p. 1643–1652.

34. Schnoegl, D., et al., AP-1 transcription factors in cytotoxic lymphocyte development and antitumor immunity. Current Opinion in Immunology, 2023. 85: p. 102397.

35. Lin, S.-Y., et al., Single-cell RNA sequencing reveals the heterogeneity and regulatory modules of cell-specific RNA-binding proteins in patients with systemic lupus erythematosus. Biochemistry and Biophysics Reports, 2025. 42: p. 101977.

36. Qiu, X., et al., Reversed graph embedding resolves complex single-cell trajectories. Nature methods, 2017. 14(10): p. 979–982.

37. Cao, E.Y., J.F. Ouyang, and O.J. Rackham, GeneSwitches: ordering gene expression and functional events in single-cell experiments. Bioinformatics, 2020. 36(10): p. 3273–3275.

38. Aibar, S., et al., SCENIC: single-cell regulatory network inference and clustering. Nature methods, 2017. 14(11): p. 1083–1086.

39. Gorji, A.E., et al., Investigation of systemic lupus erythematosus (SLE) with integrating transcriptomics and genome wide association information. Gene, 2019. 706: p. 181–187.

40. Richter, P., et al., Cytokines in systemic lupus erythematosus—focus on TNF-α and IL-International Journal of Molecular Sciences, 2023. 24(19): p. 14413.

41. Xu, H., et al., Integrated bioinformatics and validation reveal TMEM45A in systemic lupus erythematosus regulating atrial fibrosis in atrial fibrillation. Molecular Medicine, 2025. 31(1): p. 104.

42. Lin, C.-H., et al., IκBα promoter polymorphisms in patients with systemic lupus erythematosus. Journal of Clinical Immunology, 2008. 28(3): p. 207–213.

43. Curran, T. and B.R. Franza Jr, Fos and Jun: the AP-1 connection. Cell, 1988. 55(3): p. 395–397.

44. Adrianto, I., et al., Association of a functional variant downstream of TNFAIP3 with systemic lupus erythematosus. Nature genetics, 2011. 43(3): p. 253–258.

45. Yang, F., J. Lin, and W. Chen, Post-translational modifications in T cells in systemic erythematosus lupus. Rheumatology, 2021. 60(6): p. 2502–2516.

46. Huangfu, L., et al., The IL-17 family in diseases: from bench to bedside. Signal transduction and targeted therapy, 2023. 8(1): p. 402.

47. Ghorbaninezhad, F., et al., Tumor necrosis factor-α in systemic lupus erythematosus: Structure, function and therapeutic implications. International Journal of Molecular Medicine, 2022. 49(4): p. 43.

48. Moulton, V.R. and G.C. Tsokos, Abnormalities of T cell signaling in systemic lupus erythematosus. Arthritis research & therapy, 2011. 13(2): p. 207.

49. Wolf, F.A., P. Angerer, and F.J. Theis, SCANPY: large-scale single-cell gene expression data analysis. Genome biology, 2018. 19(1): p. 15.

50. Virshup, I., et al., anndata: Annotated data. BioRxiv, 2021: p. 2021.12. 16.473007.

51. Wolock, S.L., R. Lopez, and A.M. Klein, Scrublet: computational identification of cell doublets in single-cell transcriptomic data. Cell systems, 2019. 8(4): p. 281–291. e9.

52. Fang, Z., X. Liu, and G. Peltz, GSEApy: a comprehensive package for performing gene set enrichment analysis in Python. Bioinformatics, 2023. 39(1): p. btac757.

53. Kanehisa, M. The KEGG database. in ‘In silico’simulation of biological processes: Novartis Foundation Symposium 247. 2002. Wiley Online Library.

54. Jin, S., et al., Inference and analysis of cell-cell communication using CellChat. Nature communications, 2021. 12(1): p. 1088.

